# Mechanism of tension propagation in cell membranes

**DOI:** 10.1101/2023.03.22.533804

**Authors:** Avishai Barnoy, Andrey K. Tsaturyan, Michael M. Kozlov

## Abstract

The propagation of the membrane tension perturbations is a, potentially, essential mechanism of the mechanical signal transduction along surfaces of live cells. The efficiency of this process is determined by the propagation speed, which turned to be a hot and a controversial topic of the Cell Biophysics. In a stark contrast to the earlier results and expectations, the recent studies in several cell types revealed a wide range of the tension propagation speeds beginning from the strikingly low ones challenging the significance of the process and up to relatively high biologically relevant rates. The previously suggested models of the tension propagation have been based on assuming an unrealistic softness of the membranes for the stretching-compression deformations, which challenges the model ability to account for the observations. Here, we consider a different physics of the generation and the propagation of tension perturbations in cell membranes. We propose the tension to be controlled by an intra-cellular pressure and the propagation of the tension perturbations to be mediated by a membrane area redistribution between compartments, to which cell membranes are divided by the proteinic barriers, according to the picket-fence model. Using the established elastic features of cell membranes including their effective non-stretchability, this mechanism quantitatively accounts for the slowness of the propagation process and gives a natural explanation of the wide range of the observed propagation speeds. The model predictions are amenable to a direct experimental verification by controlled osmotic pressure variations.

## Introduction

The membrane tension and the speeds of propagation of its perturbations along cell membranes have recently become one of the hot and controversial topics discussed in the cell biological literature(1-4). Our goal here is to propose a physical mechanism for the propagation of the tension perturbations, which resolves the current controversies and is amenable to straightforward experimental verifications.

From the physics perspective, the biological membranes are few nanometer-thick continuous films exhibiting an in-plane fluidity and an elastic resistance to the deformations of bending and stretching-compression(5, 6). The fluidity and elasticity originate from those of lipid bilayers, which serve as the membrane structural matrices(7) and are mechanically stabilized by the powerful hydrophobic effect(8). Since the thickness of a biological membrane is, typically, several orders of magnitude smaller than the membrane in-plane dimensions, the membranes are commonly described as 2D elastic fluids (5).

The membrane tension (5, 6, 9) is the stress emerging in response to the application to the membrane of stretching (leading to positive tensions) or compressing (leading to negative tensions) forces. The membrane tension is in many respects a 2D analogue of the pressure in 3D fluids. Based on this analogy, a tension perturbation generated at some membrane location by local forces has been expected to rapidly propagate along the whole membrane plane(10). Due to this straightforward reasoning, the tension has been considered an efficient mediator of cross-talks between mechanical events happening at different locations along the membrane (11, 12). Specifically, the tension in the external cell membrane, the plasma membrane, was proposed to coordinate the cell surface processes such as the endo- and exocytosis (13-16), the cell movement and shape transformations (10, 17-21), the cell signalling (22, 23), the axon growth and branching (24).

The efficiency and, hence, the biological relevance of the tension as a signal transducer critically depend on the speed with which a local tension variation spreads along the membrane. Initially, the propagation of a tension perturbation has been considered instantaneous compared to the biologically relevant times of seconds(9-11, 17, 18, 21, 25-30). Yet, this notion has been challenged by the study of the propagation dynamics of the tension perturbations in the plasma membranes of live HeLa cells(1). The experiments were performed by the dual-tether assay, in which the background tension was perturbed by pulling of a tether from the cell membrane and recorded by a second tether located at a desired distance from the first one(1). No measurable tension propagation was registered within a time span of about ten minutes between the tethers pulled out of the regular regions of the plasma membrane and separated by the 5 - 10 μm distances. The qualitatively similar results were obtained for several types of non-motile cells(1). A relatively fast spreading of the tension, happening within a timescale of a few seconds, was observed only between tethers pulled out of a cell bleb (1), which is a spherical outgrowth of the plasma membrane devoid of the sub-membrane layer of the cortical cytoskeleton (31).

Studies aiming at an exploration and a verification of the phenomenon of the ultra-slow spreading of the tension perturbations resulted in a complex picture. The speed of the process varied depending on the cell type, the region of the plasma membrane, and the way of the force application to the cell surface (3, 4, 24) over orders of magnitude wide range. In a stark contrast to the results of (1), application of the dual-tether assay to the goldfish retinal bipolar neurons revealed the tension perturbations to be rapidly transmitted within a few seconds to distances of 4 to 10 μm (4). This result was obtained for the membranes of both the presynaptic terminals and the soma of these neurons. Testing in the same study the tension propagation in the membranes of the neuroendocrine chromaffin cells did not reveal any measurable tension transmission to the 6 – 12 μm distances within several minutes(4), similar to the previous data on HeLa cells(1). The large speed of about 20 μm/s was measured for the transmission of the membrane tension perturbations in the axons of the rat hippocampal neurons(24). Yet, in contrast to the axons, no tension propagation within the relevant timescale was registered for the dendrites of the same cells, consistent again with the previous results(1). A relatively rapid propagation within several seconds to about 10 μm distances was observed for the membrane tension perturbations produced by the activation of the localized actin-driven membrane protrusions in the neutrophil-like cells (3). Strikingly, the dual-tether assay in the same cells revealed no membrane tension spreading even if the tethers were separated by less than 2μm distances (3). Summarizing, the rates of the propagation of the tension perturbations measured in the plasma membranes of the diverse cells varied in the range between more than 10μm/s and less than 1μm/min.

An understanding of the physical mechanism behind the extreme slowness of the tension propagation in the particular cell membranes, and of a, generally, wide variation range of the observed speeds of this process poses a challenge. Indeed, if a membrane was naively considered as a simple continuous elastic sheet suspended in an aqueous environment, the tension perturbations were expected to propagate with a speed of sound, which for lipid bilayers was estimated to be in the range 0.1-1.0 m/s (32), hence, exceeding by several orders of magnitude the highest observed rates of about 10μm/s. Thus, the observed rates of the tension transmission and, especially, the ultra-slow ones, must result from the membrane interactions with other cell structures. It has been suggested and experimentally substantiated that this phenomenon is governed by a mechanical cross-talk between the membrane and the underlying cortical cytoskeleton, and mediated by the proteins spanning the lipid bilayer and linking the membrane to the cortical layer(1-3, 24).

To put this hypothesis on a quantitative basis and relate it to the measurable material parameters of the lipid membranes, a physical model was proposed in which, according to the classical view, the perturbations of the membrane tension were related to the variations of the membrane in-plane density. The speed of the propagation of the tension perturbations was assumed to be restricted by the viscous friction within the membrane flow induced by the spreading of the density variations. The cortex-anchored trans-membrane proteins were considered to play a role of the two-dimensional obstacles resisting the membrane flow and generating the major friction ^1,3,24^.

While these models offered a qualitative explanation of the phenomenon of a slow, as compared to the sound waves, propagation of the membrane tension perturbations, their ability to account for the observed speeds of this process remained debatable. Specifically, because of the postulated direct relationship between the viscous membrane flow and the dynamic variations of the membrane density, these models predicted the speed of the tension propagation to be proportional to the local stretching modulus of the membrane ^1-3^. The values of this modulus were established to be of the order of 10^5^pN/mm ^6^. To get a quantitative agreement between the theory’s predictions and the measured speeds of the tension propagation, the previous models used much lower values of the stretching modulus, such as 40pN/μm ^1,2^ and 100pN/μm ^3^. The former value originated from the measurements of the elastic deformations of neutrophils and characterized the rigidity of the cytoskeletal networks of these cells rather than that of the cell membranes, as was explicitly stated in the original articles ^33,34^. The latter value was taken from^9^, where it was attributed to the membrane tension rather than the stretching modulus. This means that the membranes have been considered extremely soft as compared to the real cell membranes and lipid bilayers. If the quantitative estimations of^1-3^ used the proper values of the stretching modulus, the predicted speeds of the tension propagation would exceed the experimentally observed values by at least three orders of magnitude.

Here we propose a different mechanism for the generation and the propagation of the tension perturbations, which does not rely on the membrane density variations and accounts for both the low values and the large variation range of the propagation speed. The model uses the established values of the membrane visco-elastic parameters, and, specifically, considers the membrane to be non-stretchable due to the discussed above high value of the stretching modulus. The mechanism is based on the picket-fence model of cell membranes^35^, which proposes the membranes to be subdivided into about 30-230nm-large compartments by a network of the trans-membrane proteinic fences-boundaries. These boundaries are resistant to the 2D diffusion of proteins and lipids and are linked to the cortical cytoskeleton^36,37^. The major idea of our model is that the membrane tension is produced within each compartment by the intracellular pressure, which pushes the compartment membrane and sculpts it into a nearly dome-like shape. A tension perturbation is generated, initially, within one compartment by reducing the excess of the compartment membrane area by, for example, pulling a membrane tether, which results in an increase in the compartment’s membrane tension. This gives rise to a membrane flow from the surrounding membrane compartments into the perturbed one, which, in turn, results in the perturbation propagation along the whole system. The speed of this propagation is largely set by the intra-cellular pressure.

Using the biologically plausible values of the system parameters, we demonstrate that the small propagation speeds of the tension perturbations result from the low intra-cellular pressures characterizing, according to the literature, the cell interior upon regular conditions. Mild variations of the intra-cellular pressure due to, for example, osmotic effects, easily explain the broad range of the observed speeds. The critical role of the intracellular pressure in the membrane tension dynamics must be easily amenable to experimental verifications.

## Model background

We consider a cell plasma membrane attached to an underlying layer of cytoskeletal cortex(33) referred to as the membrane skeleton (34, 35) (Fig.1). The membrane is regarded as a continuous fluid elastic film, which can undergo an in-plane flow, sustain a hydrostatic pressure difference of, for example, an osmotic origin, and develop a membrane tension(6). The membrane skeleton is a network of contractile fibers representing the actin-myosin complexes(33). Because of the tens of nanometer large spaces between the constituent fibers, the membrane skeleton is fully permeable to water and, therefore, does not directly hold the hydrostatic pressure. The fibers develop contractile forces described as the cortical tension, which drives the network shrinkage (36). The cortical tension can change upon variable intra-cellular conditions and adopts values of few tenths of mN/m during the interphase, and several mN/m during the metaphase (37).

As a base for the mechanical model of the membrane-cortex system, we consider the two previously elaborated features of cell plasma membranes: the membrane compartmentalization by the cortex described by the picket-fence model (34, 38-40); and the mechanical interplay between the excess of the intracellular hydrostatic pressure and the cortical tension (37, 41).

According to the picket-fence model, the membrane is subdivided into compartments separated by the fences-boundaries, which are formed by the rows of integral picket-proteins, span the membrane thickness, and are anchored to the membrane skeleton (34). These fences-boundaries impede the in-plane diffusion of the lipid molecules in the two membrane leaflets(42, 43). Since the picket-proteins must represent obstacles also for the in-plane lipid flow(44), we consider the boundaries to be characterized by a certain hydrodynamic permeability for the membrane flow. The in-plane dimensions of the membrane compartments vary in the range between tens to hundreds of nanometers(35, 45).

The existence and extent of an excess of the intracellular hydrostatic pressure as compared to the pressure in the extracellular medium(41), referred below to as the intracellular pressure, was revealed and quantified by the micromechanical experiments (37, 41). The intracellular pressure originates from the trans-membrane osmotic gradients (41) and can be controlled by changing the cell volume or the amount of the osmolytes (37). The intracellular pressure varies between few tens Pa and few thousands Pa depending on the cell type and the stage of the cell cycle(46).

The intracellular pressure and the cortical tension counteract and their values are interdependent, as evidenced by the experiments in which stimulations of a contraction of the actomyosin cytoskeleton increased whereas a disruption of the actomyosin activity reduced the intracellular pressure (41). Moreover, the cortical tension and the intracellular pressure mutually equilibrate since their values could be related through the Laplace equation involving the curvature of the apparent cell surface (37). Finally, the variations of the intracellular pressure were accompanied by the substantial (10-40%) changes of the volumes and, hence, the apparent surface areas of the rounded cells (41). Taking into account that the membrane is, practically, non-stretchable, the large surface area variations of the effective sphere must be due to a contraction of the cortical membrane skeleton (47, 48), while the membrane is folded and serves as an area reservoir by smoothening its folds during the stretching of the apparent surface of a cell(49).

## Model

Based on the above concepts, we formulate the following model for the tension generation and propagation within the membrane coupled to the cortical membrane skeleton.

### The system structure and the physical properties

- The non-stretchable fluid membrane of a given area is attached to the underlying membrane skeleton by the rows of the trans-membrane proteins, which divide the membrane into the identical compartments and serve as the compartment boundaries (Fig. 1A).

**Fig. 1.**
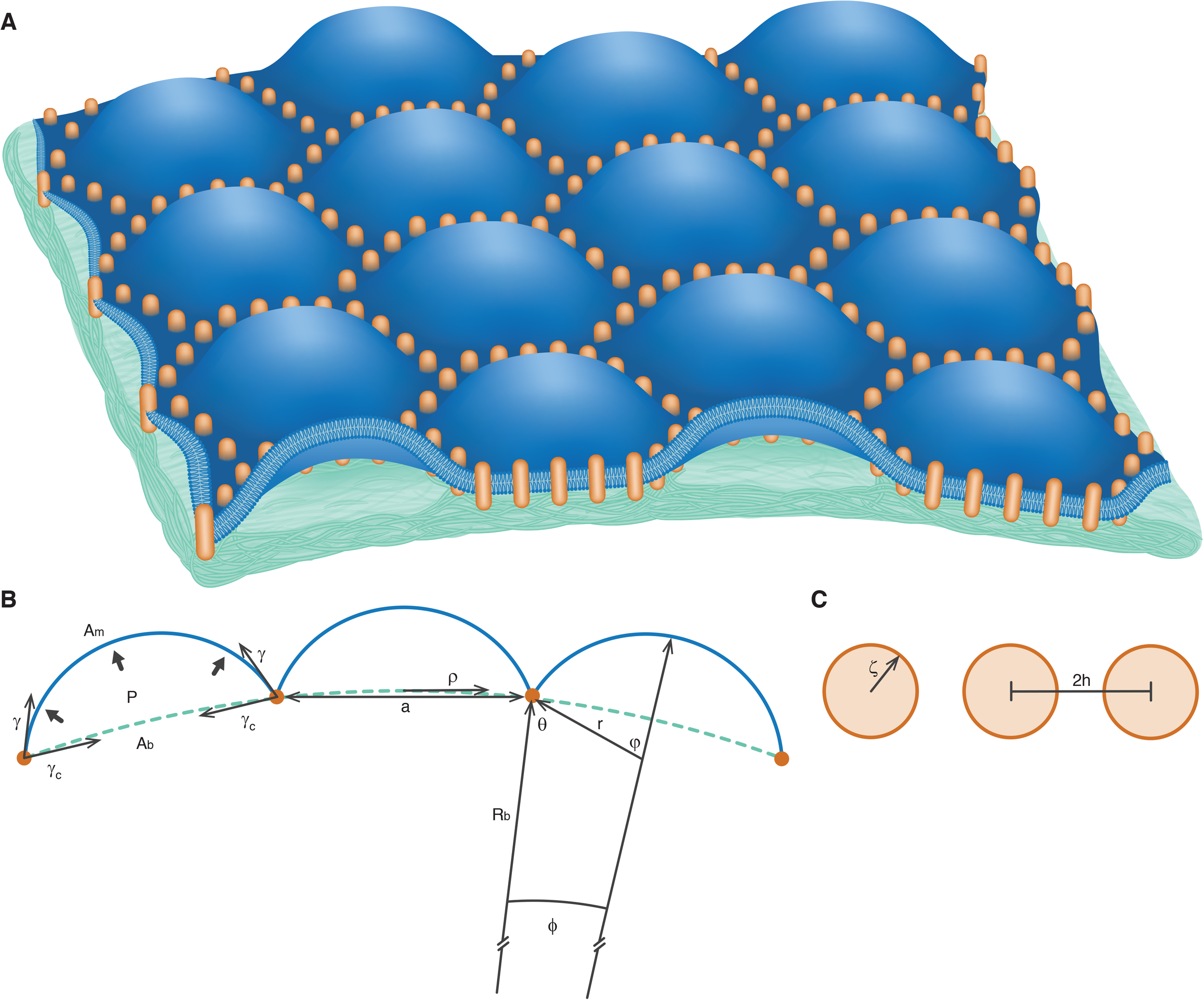
The illustration of the model and the notations. A. The membrane is connected to the membrane skeleton by rows of transmembrane picket-proteins, according to the picket-fence model (34). The protein rows subdivide the membrane into compartments and serve as hydrodynamic barriers for the membrane flow. The membrane within each compartment bulges outwards because of a contraction of the membrane skeleton. (B) A schematic representation of the cross-sections of the dome-like shapes of the membrane compartments and the notations. (C) A schematic representation of an element of the compartment boundary and the notations. The compartment boundaries form a 2D network, which lies in a plane referred below to as the membrane base plane. We assume the base plane to have a shape of a spherical segment with a curvature radius, *R*_*b*_ (Fig. 1B). The area occupied by one compartment in the base plane is referred to as the compartment base area, *A*_*b*_, which corresponds to the linear dimension, 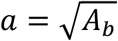 (Fig. 1B). We assume the radius of curvature of the base plane to largely exceed the compartment size, *R*_*b*_ ≫ *a*, and all the compartments to have the same size, *a*. The membrane area enclosed within one compartment, *A*_*m*_, is referred below to as the compartment membrane area. Because of the actomyosin contractility of the underlying membrane skeleton(36), the boundary of each compartment is subject to the cortical tension, *γ*_*C*_, which acts to shrink the compartment base area, *A*_*b*_ (Fig. 1B). This results in an excess of the compartment membrane over the base area, *A*_*m*_ *− A*_*b*_, and a related bulging of the compartment membrane outwards of the base plane (Fig.1A, B)).
- The system is subject to an intracellular pressure, *P* (Fig. 1B), which acts to increase the volume bounded by the cell membrane and, hence, to stretch the base plane, in general, and the base area on each compartment, *A*_*b*_, in particular. This stretching is counteracted by the cortical tension, *γ*_*C*_, acting to reduce *A*_*b*_. We assume the cortical tension to be sufficiently large so that the excess area, *A*_*m*_ − *A*_*b*_, does not vanish and the related membrane bulging persists.
- Since the membrane skeleton is permeable to water, the intracellular pressure, *P*, pushes the membrane within each compartment in the outward direction and generates a membrane tension, *γ*. Finding *γ* as a function of *P* includes as an intermediate step a computation of the membrane shape within the compartment. Here we assume, for simplicity, that the compartment boundary is circular, and shape of the compartment membrane is that of a spherical segment referred below to as the spherical dome (Fig. 1B). In this case the membrane tension is related to the intracellular pressure by Laplace equation

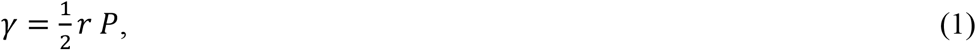

where *r* is the radius of the spherical dome approximating the compartment membrane shape (Fig. 1B). The membrane tension, *γ*, is constant within one compartment, but can be different for different compartments.

In (SI E) we describe the results of a more detailed analysis accounting for the boundary effects and demonstrate that the predictions of the model remain qualitatively unchanged.

- If the membrane tension, *γ*, is different in two adjacent compartments, the membrane tends to flow across the compartment boundary from the compartment with a lower to that with a higher *γ*. The compartment boundaries resist this flow, which we quantify by the hydrodynamic permeability of the boundary, *λ*, such that the flux of the membrane area through the unit length of the boundary, *q*, is related to the trans-boundary difference of the membrane tension, Δ*γ*, by

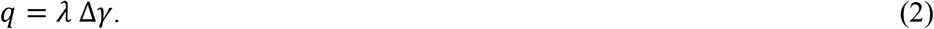

The boundary permeability, *λ*, is determined by the hydrodynamic interaction between the trans-membrane picket-proteins building up the boundary and the membrane flow (44). We model these proteins as cylindrical rods (Fig. 1A) with cross-sectional radius, ζ (Fig. 1C). The rods are oriented perpendicularly to the membrane plane and separated by an in-plane distance, 2*h*, between the centers of their cross-sections (Fig. 1C). Based on the results of (50, 51), the boundary permeability is given by (for the details see SI A)

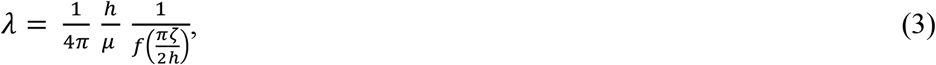

where *μm* is the 2D dynamic viscosity of the membrane and the non-linear function 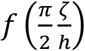 presented in (SI A).

### The mechanism of the induction and the propagation of the membrane tension perturbation

- Initially, the system is in a mechanical equilibrium, which requires a mutual balancing between the intracellular pressure, *P*, the membrane tension, *γ*, and the cortical tension, *γ*_*C*_. For the relevant case of *R*_*b*_ ≫ *r*, this equilibrium is determined by (for a derivation see SI B)

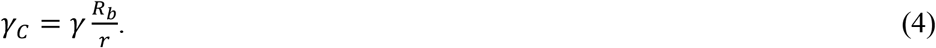

In addition, in equilibrium the inter-compartment membrane flow must vanish meaning that the membrane tension, *γ*, must be equal in all the compartments.
- The equilibrium is perturbed by an instantaneous extraction of some amount of the membrane area from one of the compartments, for example, by pulling out of it a membrane tether(1, 24). As it will be shown below, the decrease of the compartment area results in an increase of the pressure-generated membrane tension within the perturbed compartment. The resulting disbalance of the tension between the perturbed and the adjacent compartments induces an inter-compartment membrane flow through the compartment boundaries. This results in the propagation throughout the system of the variations of the membrane area and, hence, of the tension perturbation.

The compartment base area, *A*_*b*_, will be assumed to remain constant during the propagation process since it is determined by the constant values of the cortical tension, *γ*_*C*_, and the intracellular pressure, *P*.

This propagation ceases when the induced perturbation is compensated by the membrane area redistribution throughout the system and the tension is equalized again in all the compartments.

## Main Equations

Our goal here is to derive the equations for the time evolution and the space distribution of the membrane tension, *γ*, and the compartment membrane area, *A*_*m*_. We assume, for simplicity, the system to be axially symmetric, and describe the position by a polar coordinate, *ρ*, measured along the base plane. The latter is considered flat due to the largeness of its radius of curvature compared with the compartment size, *R*_*b*_ ≫ *a*.

As a parameter characterizing the compartment membrane area, we use its relative excess,

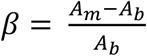 referred below to for brevity as the excess area. In the following we consider flat domes, *β* ≪ 1.

Initially, all the compartments have the same value, *β*_0_, of the excess area. The perturbation of *β* is induced at the time point, *t* = 0, in the central compartment with a midpoint at *ρ* = 0, and radially spreads into the surrounding compartments. This is accompanied by a change of the membrane tension, *γ*, in the perturbed and then in the other compartments. While, because of the finite compartment size, both *β* and *γ* are discretely distributed along the base plane, we will use the continuous approximation, in which the time evolution of the system is described by the continuous functions *β(t, ρ*) and *γ*(*t, ρ*) (for the details see (SI C)). The continuous description is valid for sufficiently large distances, *ρ* ≫ *a* (SI C).

The equations for *β*(*t, ρ*) and *γ*(*t, ρ*) can be obtained by considering the dynamic changes of the compartment membrane areas, *A*_*m*_, due to the inter-compartment membrane fluxes (Eq.2), and by accounting for the mechanical interplay between the tension and the intracellular pressure.

The area dynamics provides a partial differential equation (SI C)

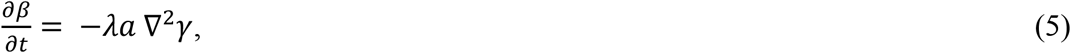

where ∇^2^ is the 2D Laplace operator in the base plane coordinates.

The force balance along with a geometrical relationship between the membrane dome radius,

*r*, and the compartment membrane area, *A*_*m*_, result in the relationship between *β* and *γ* (for the details see (SI D))

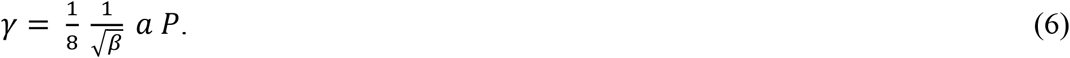

The (Eqs. 5,6) can be combined in a non-linear partial differential equation either for the area excess, *β*,

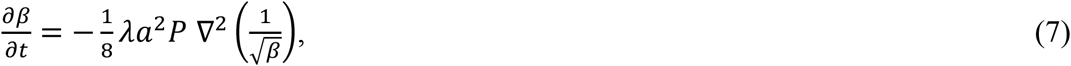

or for the membrane tension, *γ*,

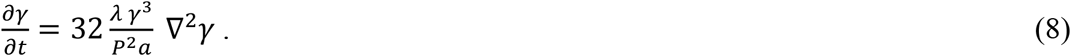

Solution of (Eq.7) or (Eq.8) requires an appropriate initial condition.

For the following, it is convenient to express the perturbations through a common function, ε(*t, ρ*), referred below to as the perturbation function. For this purpose, we present the excess area in the form

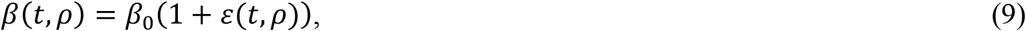

where *β*_0_ is the initial excess area, and

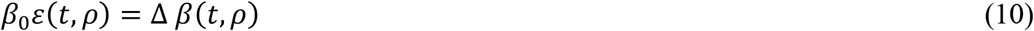

is the excess area perturbation. The membrane tension *γ*(*t, ρ*) and its perturbation, Δ*γ*(*t, ρ*) can be determined by using (Eq. 6) together with (Eq. 9).

An exact initial condition for the perturbation function would be a step-like function, ε(0, *ρ*), which adopts a constant value, ε_0_, within the central membrane compartment, ε(0, *ρ*) = ε_0_ for 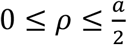, and vanishes everywhere else, *ε*(0, *ρ*) for 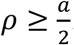. Yet, instead of this, we will use Gaussian distribution with a standard deviation equal to the compartment half-size,

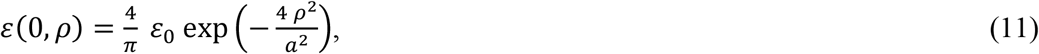

which, on one hand, greatly facilitates the calculations and, on the other, provides all the qualitative features of the system. Since within the considered scenario, the perturbation reduces the area of the central compartment, ε_0_ < 0.

## Results

For understanding the system dynamics, we consider the case of small perturbations of the excess area, ∆*β*(*t, ρ*) ≪ *β*_0_, described by small absolute values of the perturbation function,

|ε(*t, ρ*)| ≪ 1, which corresponds to, |ε_0_| ≪ 1.

Using (Eqs.7,9) and retaining only the contributions linear in ε(*t, ρ*) we obtain for the perturbation function,

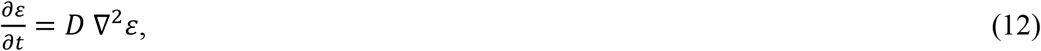

which has the form of a diffusion equation with a diffusion coefficient

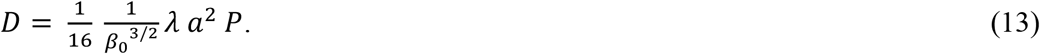

The solution of (Eq.12) accounting for the initial condition (Eq.11) is (for the details see (SI E))

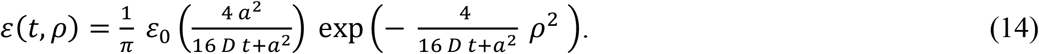

For times, 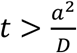, it can be presented in the familiar form describing diffusion in an infinite

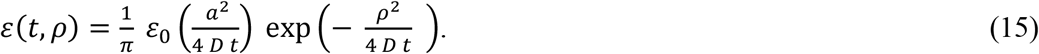

The excess area *β*(*t, ρ*), and its perturbation, ∆*β*(*t, ρ*), are expressed through ε(*t, ρ*) by the relationships (Eq.9) and (Eq.10), respectively.

To obtain the expressions for the membrane tension, *γ*(*t, ρ*), and its perturbation, ∆*γ*(*t, ρ*), we account in the general expression (Eq.6) only for the first order contributions in ε(*t, ρ*), and obtain

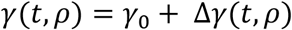

where

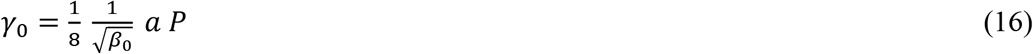

is the initial background tension and

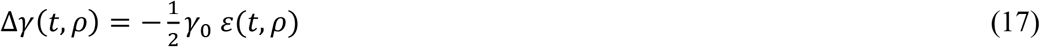

is the small tension perturbation, ∆*γ*(*t, ρ*) ≪ *γ*_0_.

### Quantitative predictions and comparisons with observations

To compare the quantitative predictions of the model with the observations, we use specific parameter values. We assume the initial excess area to be *β*_0_ = 0.1. Because the typical size of the membrane compartments varies in the range 30 – 230 nm (34, 53) we use a mid-value of *a* = 100*nm*. To estimate the 2D hydrodynamic permeability, *λ*, of the proteinic fences-barriers (Fig.1C) we assume the in-plane cross-sectional radius of a trans-membrane picket-protein to be, ζ = 1*nm*, the half-distance between the adjacent picket-proteins to be, *h* = 2.5*nm*, and the 2D membrane viscosity to have a value 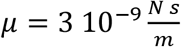. Using (Eq. 3) (see (SI A) and (51)) we obtain 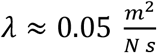. We use the two specific values of the intracellular pressure, *P* = 40 pa and *P* = 400 pa, which were measured during, respectively, the interphase and metaphase of HeLa cells (37). Based on (Eq.16), these pressure values correspond to the background membrane tension, *γ*_0_, of, respectively, 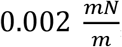 and 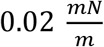. The two values of the tension belong to the tension range measure in live cells (32). Using (Eq.13) we estimate the diffusion coefficient, *D*, to be equal 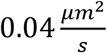 for *P* = 40 pa and 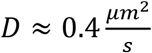 for *P* = 400 pa.

The time evolution of the perturbation function, ε(*t, ρ*), is illustrated in (Fig. 2). The propagation of the perturbation is described by widening and flattening of the perturbation function with time (Fig. 2).

**Fig. 2.**
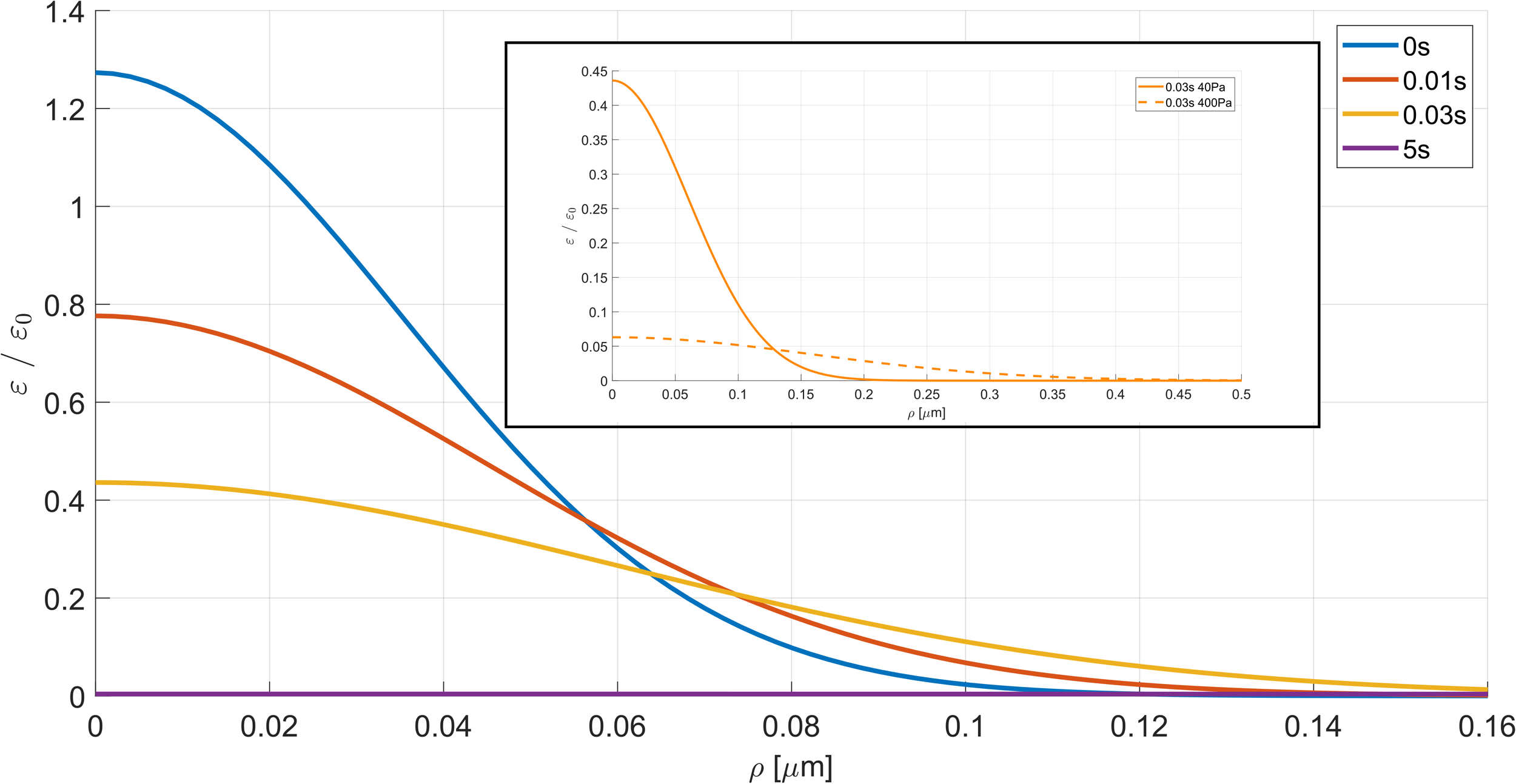
The time evolution of the perturbation function for 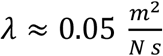 *a* = 100 *nm*, and *P* = 400 pa (dashed lines) or *P* = 40 pa (solid lines).

The propagation rate of the perturbations can be illustrated by the time dependence of ε, at three different locations as presented in (Fig. 3). For each location, *ρ*, the perturbation function reaches a maximum, ε^∗^, at a certain time point, *t*^∗^, and then decays. The dependence *t*^∗^(*ρ*) following from (Fig. 3) is presented in (Fig. 4) and describes the propagation pace.

**Fig. 3.**
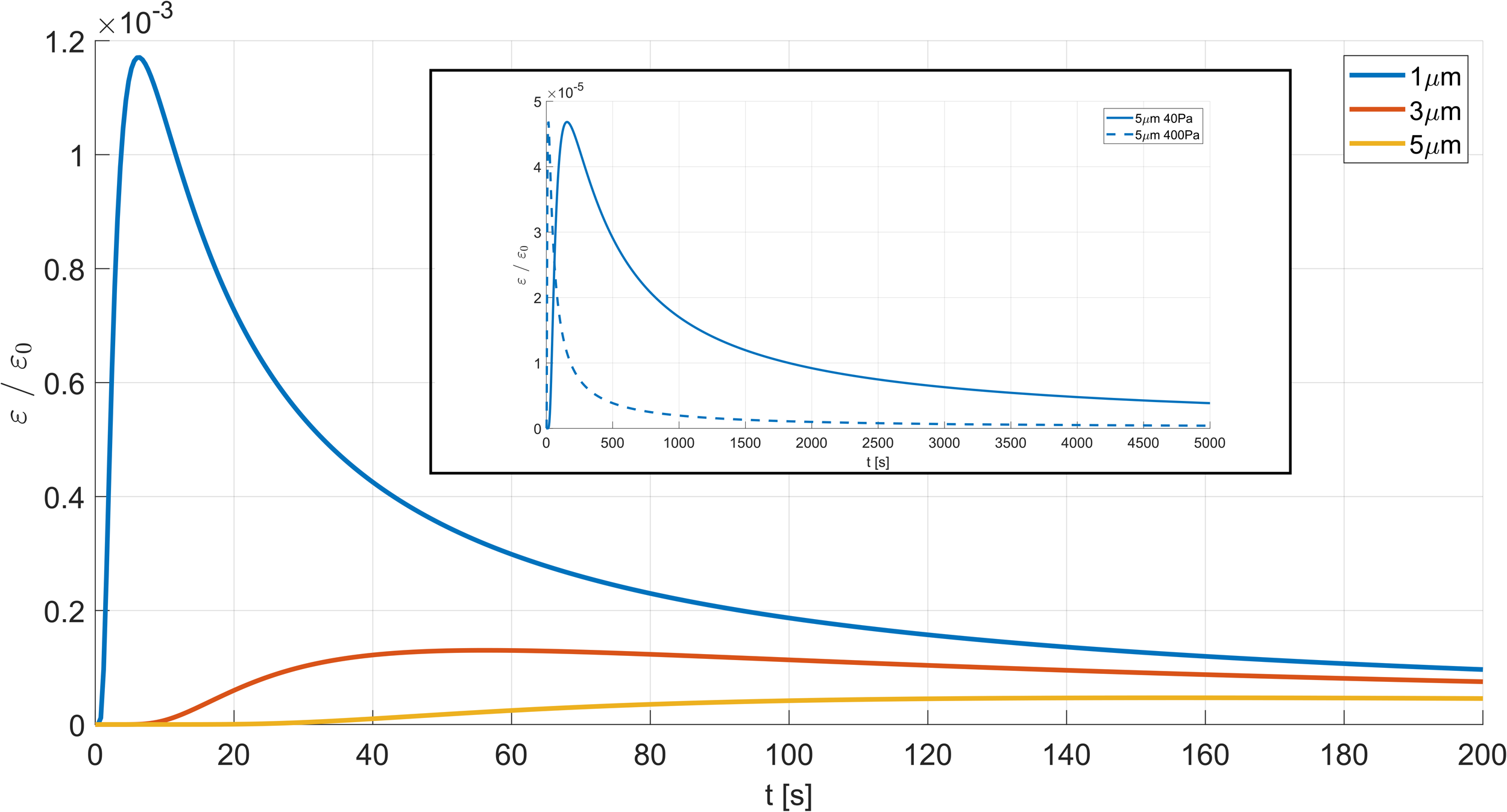
The time evolution of the perturbation function for different distances from the initially perturbed compartment. The parameter values: *β*_0_ = 0.1, 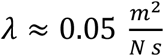, *a* = 100 *nm, P* = 400 pa for the dashed lines, *P* = 40 pa for the solid lines.

**Fig. 4.**
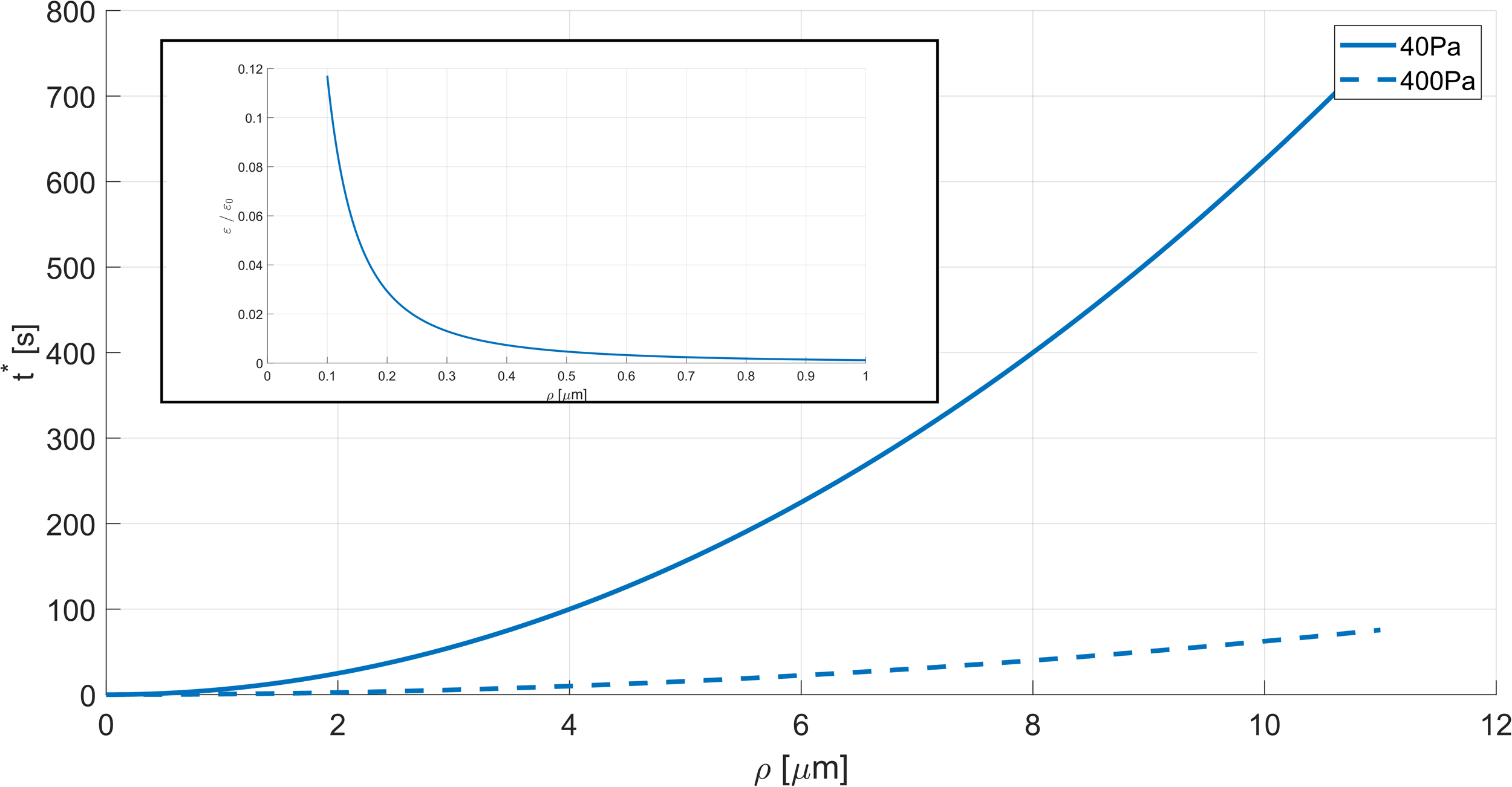
The time of reaching the maximal perturbation as a function of the distance from the initially perturbed compartment. Inset: the maximal tension perturbation. The parameter values: *β*_0_ = 0.1, 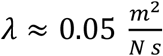, *a* = 100 *nm, P* = 400 pa for the dashed lines, *P* = 40 pa for the solid lines.

An approximate form of the function *t*^∗^(*ρ*) follows from (Eq. 14) and is given for *ρ* ≫ *a* by

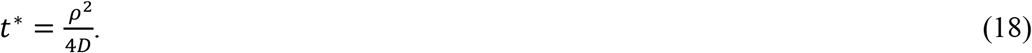

According to (Eq. 18), the time of the perturbation propagation to the experimentally relevant distance of, *ρ* = 10*μm* (1, 3), has the value of about *t*^∗^ = 10 *min* for the diffusion coefficient 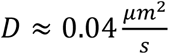 corresponding to a regular pressure of *P* = 40 pa and the above values of the structural parameters. This agrees with a slow propagation of the membrane tension perturbations, as observed in the tether pulling experiments (1, 3).

A propagation time, *t*^∗^, of few seconds, as observed in (3) upon a local stimulation of the actin polymerization in the goldfish(4) and hippocampal(24)neurons can be easily explained by a larger intracellular pressure, *P*. Specifically, for the above used parameter values, the time, *t*^∗^ ≈ 1*s*, for the distance, *ρ* = 10*μm*, requires an intracellular pressure, *P*, produced by a modest trans-membrane osmotic gradient of about 10mM.

Importantly, as it follows from (Eq. 13), higher rates of propagations may result also from larger values of the compartment size, *a*, and of the hydrodynamic permeability of the compartment boundaries, *λ*, and from smaller values of the initial excess area, *β*_0_ This is in part illustrated in (Fig. 5), which presents the dependence of the diffusion coefficient, *D*, on *β*_0_ and on the fraction of the compartments boundary length occupied by the picket-proteins.

**Fig. 5.**
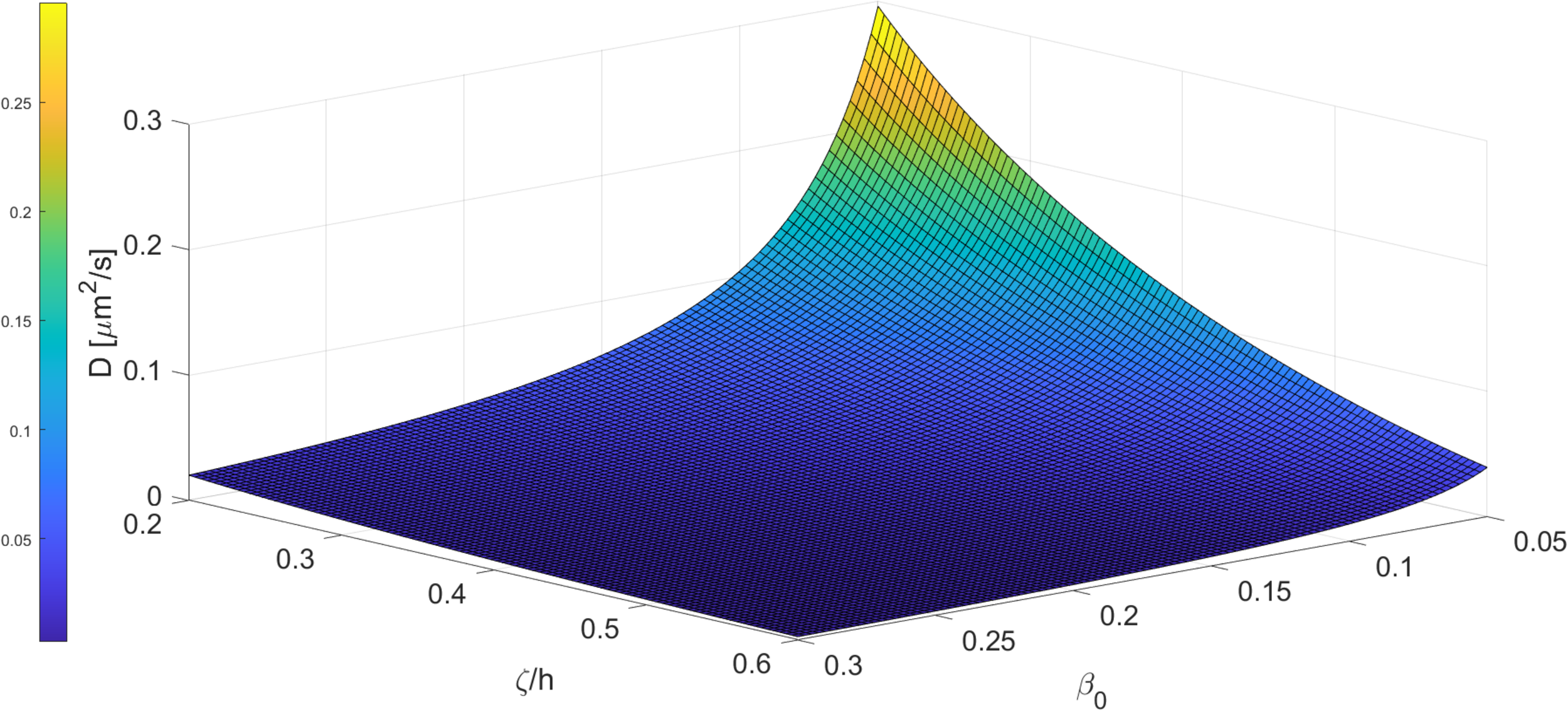
The dependence of the diffusion coefficient, *D*, on the initial excess area, *β*_0_, and density of the picket proteins at the barrier line between the compartments, ζ/*h*. Select all

Finally, the maximal value of the tension perturbation, ε^∗^, emerging at a distance, *ρ*, from the center at the time point, *t*^∗^(*ρ*), decays with the distance, *ρ*, according to

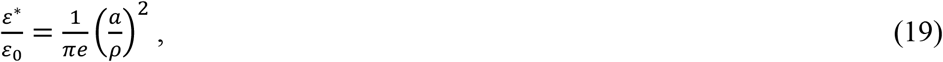

as illustrated in (Fig. 4 inset).

## Discussion

The propagation rate of membrane tension perturbations is a hot and a controversial topic of the Cell Biophysics. Initially, local variations of the membrane tension have been assumed to spread instantaneously over cell plasma membranes, hence, serving as efficient transducers of the mechanical signals along the cell surfaces. Yet, recent experiments revealed a strikingly large variation range of the tension propagation speeds, which encompasses, depending on the cell type and the cell region, very slow speeds of less than 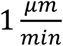 and relatively large ones exceeding 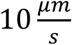 (1, 3, 4, 24). The physical mechanisms accounting for these observations have remained debatable.

Here we propose a mechanism of the tension propagation, which naturally explains the whole spectrum of the observations. The chief premise of the model is a crucial role played by the subdivision of the membrane into about 100 nm large compartments by the fences of the trans-membrane picket-proteins attached to the sub-membrane cortical cytoskeleton(34).

Extending the original idea of the picket-fence model, according to which the compartment boundaries restrict the in-plane diffusion of lipids and proteins (34, 42, 43), we assume that they also serve as hydrodynamic barriers for the in-plane flow of the membrane material.

The essence of the mechanism is that the membrane tension is produced in each compartment by the intracellular pressure, which bulges the compartment membrane into a dome-like shape and stresses it. An abrupt change of the membrane area within a certain compartment by, for example, pulling out of it a membrane tether, leads to an increase of the pressure-induced tension within this compartment. This generates membrane flow from the surrounding compartments into the disturbed one, tending to compensate for the loss of the membrane area and the related tension perturbations. The rate of this flow, which depends on the intracellular pressure and the hydrodynamic permeability of the compartment boundaries, sets the propagation speed of the tension perturbation along the system.

Based on the realistic values of the system parameters and the relevant intracellular pressure values, the model explains the whole range of the experimentally observed propagation speeds. For pressures of few tens of Pa, as measured during the interphase(41), the model predicts very slow paces of tens of minutes per micron, as initially observed(1). For larger intra-cellular pressures corresponding to tens of mM osmotic differences between the intra- and extracellular volumes, the tension perturbations are predicted to cross about 10 *μm* long distances within seconds, as observed in other cell systems(3, 4, 24).

The model predictions can be directly checked by variations of the osmotic strength of the extracellular medium, leading to controllable changes of the intracellular pressure.

Importantly, besides the intracellular pressure, the predicted propagation speed of the tension perturbation substantially depends on the other system parameters, which can vary between different cell types and intracellular conditions, such as the hydrodynamic permeability of the compartment boundaries, the membrane compartment size, the difference between the compartment membrane and base areas. The propagation speed is especially sensitive to the latter parameter (Fig. 5), which is expected to depend on the degree of contractility of the cortical cytoskeleton.

An evident difference between the suggested mechanism and the earlier proposed ones(1, 3, 42) is in the dependency of the propagation speed on the local stretching rigidity of the membrane. While our model considers the membranes to be non-stretchable, as set by the experimentally established high values of the membrane stretching moduli(6, 54), the previous models were able to explain the observed values of the propagation speeds only by assuming the membranes to be few orders of magnitude softer than the real cell membranes and lipid bilayers.

The origin of this difference is in the distinction between the physical principles underlying the present and the earlier considered mechanisms. Previously, it was inferred that the essence of the tension propagation is the transmission along the membrane surface of a local variation of the membrane density related to the tension perturbation. In this mechanism, the viscous membrane flow slowing down the propagation process strongly depends on the extent of the lipid density variation. The latter can reach significant levels only for unrealistically soft membranes, whose stretching rigidity is orders of magnitude lower than that established by numerous experimental works. In contrast, in our model, the membrane flow does not originate from the lipid density variations. It is driven by the tension differences between adjacent membrane compartments. These tension differences depend on the pressure itself and the intra-compartment excesses of the membrane area and are independent of the membrane stretching rigidity.

Discussing the previously suggested mechanisms(1-3), it is important to emphasize a potentially confusing issue of the membrane rigidity for stretching. Two notions of stretching rigidity have been introduced to describe the elastic behavior of membranes. One is the local rigidity, which sets a relationship between the membrane tension and the local changes of the in-plane density of the membrane material or, equivalently, the in-plane area per a constant number of lipid molecules. The second is the effective rigidity, which relates the tension to the global area of the membrane projection on the base plane. The two rigidity types drastically differ in both the underlying physics and the numerical values. While, as mentioned above, the local rigidity determines the relationship between the local stretching strains and stresses of the membrane material, the effective stretching rigidity describes the thermodynamic work needed for smoothening of the membrane wrinkles, which can be produced by such mechanisms as the caveolae formations(55), the thermal undulations(56), and, possibly, the cytoskeleton-related micro-protrusions (49). The value of the effective stretching rigidity is set by, generally, a weak resistance of the membrane wrinkles to smoothening and, therefore, can be orders of magnitude smaller than the local stretching rigidity. It is essential to stress that the previously proposed mechanisms were explicitly based on the local rather than the effective stretching rigidity since the membrane density fluctuations are generated within and propagate along the actual membrane plane even if the latter is wrinkled. The value of the effective stretching rigidity is, therefore, irrelevant for the speed of the tension propagation.

Finally, the analysis of the proposed model includes simplifications so that its predictions are of a semi-quantitative character. The major simplifying assumption is that of a homogeneous spherical dome-like shape of the compartment membrane, which neglects the strong variations of the membrane curvature at and in the vicinity of the compartment boundary.

These variations result from the condition of a smooth membrane shape transition between adjacent compartments. Our preliminary accounting for these boundary restrictions showed that they influence the intra-compartment membrane shape, but do not qualitatively change the model predictions concerning of the dependence of the propagations rates of the tension perturbations on the intracellular pressure and the other system parameters. Additional factors neglected by our model are the possible deviations of the compartment boundaries from the circular shape and variations of the compartment sizes leading to irregularities of the compartment networks. While, according to our estimations, also these factors do not qualitatively change the model predictions, the evaluation of the related effects requires numerical simulations, which are beyond the scope of the present work.

## Supporting information

Supplemental Derivations

## Acknowledgements

We are grateful to Haim Diamant and Naomi Oppenheimer for discussions. MMK was supported by and Israel Science Foundation grant 1994/22, and holds Joseph Klafter Chair in Biophysics.

