## Supplemental Derivations for "Mechanism of tension propagation in cell membranes"

### Supplementary Information

#### A. The hydrodynamic permeability of the inter-compartment boundaries

We consider the boundary between the membrane compartments as a 2D hydrodynamic barrier formed by a row of trans-membrane proteins, which we model as cylindrical rods oriented perpendicularly to the membrane plane. The length of each rod is equal to the membrane thickness, $d$, the rod circular cross-section in the membrane plane has radius, $\zeta$, and the in-plane distance between the rod cross-section centers is $2h$ (Fig. 1C of the main text). Our goal is to obtain an expression relating the difference in membrane tension, $\Delta\gamma$, to the flux of membrane area between the membrane bulks separated by the barrier, $q$. The 2D viscosity of the membrane, $\mu$, is assumed to be expressed through the effective dynamic viscosity of the membrane material, $\eta$, by

$$\mu = \eta d. \tag{S1}$$

The problem of a 2D flow of liquid through a row of parallel cylinders oriented perpendicularly to the flow direction was previously solved [1,2]. The drag force, T, acting on a unit length of each cylinder in the flow direction is given by

$$\mathrm{T} = 8\pi\,\eta\,U\,f\left(\frac{\zeta}{h}\right), \tag{S2}$$

where $U$ is the fluid velocity for sufficiently large distances, $L$, from the cylinder row, $L \gg \zeta + h$. The function $f\left(\frac{\zeta}{h}\right)$ is presented by

$$\frac{1}{f(t)} = 1 - 2\log(2t) + \frac{2}{3}t^2 - \frac{1}{9}t^4 + \frac{8}{135}t^6 - \frac{53}{1350}t^8 + \frac{1112}{42525}t^{10} - \frac{241643}{13395375}t^{12} + \frac{18776}{1488375}t^{14} - \cdots$$

where $t = \frac{\pi\,\zeta}{2h}^2$.

Considering that the area flux through the unit length of the rod raw is equal to the fluid velocity, $q = U$, the force acting on one rod is $\tau = \mathrm{T}\,d$, and the difference in tension between the membrane bulks separated by the rod row can be presented as

$$\Delta\gamma = 2h\,\tau. \tag{S3}$$

The relationship between the tension difference and the flux, accounting for (Eq.S1), is given by

$$\Delta\gamma = 4\pi\,\frac{\mu}{h} f(t)\,q, \tag{S4}$$

which results in the expression (Eq.3) of the main text for the 2D hydrodynamic permeability of the inter-compartment boundary, $\lambda$.

#### B. The relationship between the membrane and the cortical tensions

The membrane, $\gamma$, and cortical, $\gamma_c$, tensions are related through the condition of mechanical equilibrium of a compartment boundary. Using the geometrical definitions illustrated in (Fig. 1B of the main text), the requirement of zero force acting on the boundary in the direction normal to the base plane is

$$\gamma \sin\beta = \gamma_C \sin\phi\,, \quad \text{(S5)}$$

where $\beta = \varphi - \phi$ (Fig. 1B of the main text), which results in

$$\gamma_C = \gamma(\sin\varphi\, cot\, \phi - \cos\varphi). \quad \text{(S6)}$$

Expressing the angles through the half-size of the compartment base, $\frac{a}{2}$, the radius of the compartment dome-like membrane, $r$, and the base radius, $R_b$,

$$\sin\phi = \frac{a}{2R_b}, \quad \sin\varphi = \frac{a}{2r},$$

we obtain

$$\gamma_C = \gamma\left(\frac{R_b}{r}\sqrt{1-\left(\frac{a}{2R_b}\right)^2} - \sqrt{1-\left(\frac{a}{2r}\right)^2}\right). \quad \text{(S7)}$$

Considering $\frac{a}{2R_b} \ll 1$, and $\frac{r}{R_b} \ll 1$, we obtain

$$\gamma_C = \gamma\frac{R_b}{r}. \quad \text{(S8)}$$

#### C. The equation for the compartment area change

We consider an axially symmetric distribution of the membrane compartments along the base plane with a compartment location characterized by the radial coordinate, $\rho$, of the compartment midpoint. The tension perturbation is induced in the central compartment with $\rho = 0$.

Our aim is to derive an equation for the change in time, $t$, of the membrane area, $A_m$, of an arbitrary compartment, $A_m(t,\rho)$.

We consider a ring of compartments with the midpoint coordinate, $\rho$. The total change of the area of all these compartments, $dA(t,\rho)$, within an infinitesimal time span, $dt$, is given by the difference, $Q$, of the area fluxes through the ring's upper and lower boundaries,

$$\frac{\partial A}{\partial t} = Q. \quad \text{(S9)}$$

Using (Eq. 2) of the main text for the area flux via a boundary unit length, we obtain

$$Q = 2\pi\,\lambda\left[\left(\rho + \frac{a}{2}\right)(\gamma(\rho) - \gamma(\rho + a)) - \left(\rho - \frac{a}{2}\right)(\gamma(\rho - a) - \gamma(\rho))\right], \quad \text{(S10)}$$

where $a$ is the compartment size equal all over the system, and $\gamma$ is the membrane tension. Assuming, $\rho \gg a$, presenting the tension as series in the small value, $\frac{a}{\rho}$, and retaining the contributions of the first nonvanishing order in this ratio, we obtain from (Eq.A2)

$$Q = -2\pi\,\lambda\, a^2\rho\left(\frac{\partial^2\gamma}{\partial\rho^2} + \frac{1}{\rho}\frac{\partial\gamma}{\partial\rho}\right). \quad \text{(S11)}$$

Taking into account that the number of the compartments within the ring is $2\pi\frac{\rho}{a}$, dividing the total flux difference (Eq.S11) by this number, and using (Eq.S9), we obtain for the compartment membrane area

$$\frac{\partial A_m}{\partial t} = -\lambda\, a^3\, \nabla^2\gamma, \tag{S12}$$

where $\nabla^2\gamma = \frac{1}{\rho}\left(\rho\frac{\partial\gamma}{\partial\rho}\right)$ is the radial part of the Laplace operator in cylindrical coordinated. Considering the relative excess of the membrane area, $A_m$ with respect to the compartment base area, $A_b$, defined by $\beta = \frac{A_m - A_b}{A_b}$, and accounting for $A_b = a^2$, we obtain from (Eq. S12),

$$\frac{\partial\beta}{\partial t} = -\lambda\, a\, \nabla^2\gamma. \tag{S13}$$

#### D. The relationship between the membrane tension and the intracellular pressure

Here, we obtain an expression relating the membrane tension, $\gamma$, the intracellular pressure, $P$, and the relative excess area, $\beta$, upon the assumption used in the main text, according to which the shape of the compartment membrane is that of an ideal spherical segment.

We use the geometrical relationships for the compartment membrane area, $A_m$, compartment base area, $A_b$, the membrane dome radius, $r$, and the dome angle, $\varphi$ (Fig. 1B of the main text),

$$A_m = 2\pi r^2(1 - \cos\,\varphi) \text{ and } A_b = \pi r^2(\sin\varphi)^2, \tag{S14}$$

the definition, $\beta = \frac{A_m - A_b}{A_b}$, of the excess area, and the relationship $A_b = \frac{1}{4}\pi a^2$, where $a$ is the compartment size.

Assuming $\beta \ll 1$, we obtain from (S14)

$$r = \frac{1}{4\sqrt{\beta}}\, a. \tag{S15}$$

Using the Laplace equation, (Eq.1) of the main text, gives

$$\gamma = \frac{1}{8\sqrt{\beta}}\, aP. \tag{S16}$$

#### E. Solution of the diffusion equation

To solve the diffusion equation for the perturbation with the initial condition, (Eqs.13,10) of the main text, we use the approach based on the function presentation as a Fourier integral in the spatial coordinates and solving the elementary differential equation for the time dependence of the Fourier components [3]. The solution of the 2D problem (Eq.13) is given by

$$\varepsilon(t,\rho) = \frac{1}{4\pi D t}\int \varepsilon(0,\bar{\rho})\exp\left(-\frac{(\rho-\bar{\rho})^2}{4Dt}\right) d\bar{A}, \tag{S17}$$

where the integration is performed over the area, $\bar{A}$, of the membrane plane, which is considered within our approximation to be flat.

Insertion into (Eq. S17) of the initial condition, (Eq.11), of the main text and integration results in (Eq.14) of the main text.
